# The transmembrane glycoprotein Gpnmb is required for the immune and fibrotic responses during zebrafish heart regeneration

**DOI:** 10.1101/2024.09.11.612527

**Authors:** Savita Gupta, Gursimran Kaur Bajwa, Hadil El-Sammak, Kenny Mattonet, Stefan Günther, Mario Looso, Didier Y. R. Stainier, Rubén Marín-Juez

## Abstract

Myocardial infarction occurs when coronary supply of oxygen and nutrients to part of the heart is interrupted. In contrast to adult mammals, adult zebrafish have a unique ability to regenerate their heart after cardiac injury. Several processes are involved in this regenerative response including inflammation, coronary endothelial cell proliferation and revascularization, endocardial expansion, cardiomyocyte repopulation, and transient scar formation. To identify potential regulators of zebrafish cardiac regeneration, we profiled the transcriptome of regenerating coronary endothelial cells at 7 days post cryoinjury (dpci) and observed the significant upregulation of dozens of genes including *gpnmb*. Gpnmb (glycoprotein non-metastatic melanoma protein B) is a transmembrane glycoprotein implicated in inflammation resolution and tissue regeneration. Transcriptomic profiling data of cryoinjured zebrafish hearts reveal that *gpnmb* is mostly expressed by macrophages. To investigate *gpnmb* function during zebrafish cardiac regeneration, we generated a full locus deletion (FLD) allele. We find that after cardiac cryoinjury, animals lacking *gpnmb* exhibit neutrophil retention and decreased macrophage recruitment as well as reduced myofibroblast numbers. Moreover, loss of *gpnmb* impairs coronary endothelial cell regeneration and cardiomyocyte dedifferentiation. Transcriptomic analyses of cryoinjured *gpnmb* mutant hearts identified enhanced collagen gene expression and the activation of extracellular matrix (ECM) related pathways. Furthermore, *gpnmb* mutant hearts exhibit larger fibrotic scars revealing additional defects in cardiac regeneration. Altogether, these data indicate that *gpnmb* expressing macrophages modulate inflammation and ECM deposition after cardiac cryoinjury in zebrafish and further highlight the importance of this subset of immune cells to support a regenerative response.

## 1. Introduction

Cardiovascular diseases (CVDs) are the leading cause of mortality worldwide with coronary heart disease (CHD) accounting for most of these deaths^1^. CHD is caused by the occlusion of coronaries that supply the heart muscle with nutrients and oxygen. This occlusion leads to the death of downstream tissues and results in a fibrotic scar^2^. In adult mammals, a fibrotic scar replaces the damaged tissue, impairing cardiac function and eventually leading to heart failure^3^. In contrast, adult zebrafish have a unique ability to regenerate their heart after cardiac injury^4-6^.

Following cardiac cryoinjury, the zebrafish heart undergoes a rapid and efficient repair process involving inflammation, neovascularization, and transient fibrotic tissue deposition. Eventually, the scar is cleared and cardiac tissues are regenerated^6-10^. After cardiac injury, the immune cells are the first to respond to tissue damage, rapidly recruited to the injured area along with coronary sprouting within hours after injury^9,11-13^. In this study, detailed transcriptomic profiling of coronary endothelial cells (cECs) identified *gpnmb* to be upregulated in regenerating cECs at 7 days post cryoinjury (dpci), a time-point when revascularization is vigorous and coronaries cover the injured area^9,12^.

Gpnmb (Glycoprotein non-metastatic melanoma protein B) is a type I transmembrane protein also known as Osteoactivin (OA); it contains a large extracellular domain and a short cytoplasmic domain localized at the cell surface and at phagosomal membranes^14,15^. ADAM10 is a protease that releases the soluble extracellular domain of Gpnmb, which itself induces endothelial cell migration and angiogenesis^15^. GPNMB extracellular domain (ECD) functions differently in different cell types based on its interacting receptors like Na^+^, K^+^-ATPase (NKA), Epidermal Growth Factor Receptor (EGFR), Vascular Endothelial Growth Receptor (VEGFR) and CD44^16-19^. Furthermore, GPNMB expressing macrophages play an important role in extracellular matrix (ECM) remodeling and fibroblast activation, and they also contribute to the balance between fibrosis and fibrolysis^20,21^. GPNMB is lowly expressed in healthy tissues but it is strongly induced in immune cells such as macrophages and monocytes following ischemic damage in kidney where it promotes regeneration^17,22^. GPNMB expression is not detectable in resident macrophages in healthy mouse hearts^23^. However, GPNMB is strongly upregulated by macrophages infiltrating into the heart after myocardial infarction (MI) in non-regenerative mouse models, and it is associated with adverse left ventricular remodeling in different cardiomyopathy models^24^. Yet, it remains unknown how Gpnmb regulates cardiac regeneration.

In this study, we used *gpnmb* full-locus deletion mutants and transcriptomic profiling to identify regenerative responses during zebrafish cardiac regeneration. We find that *gpnmb* is induced in macrophages after injury and that its deletion alters the inflammatory response affecting neutrophil and macrophage recruitment. Furthermore, *gpnmb* mutants exhibit myofibroblast differentiation defects and an increased fibrotic response. Our data also reveal the essential role of Gpnmb in ECM remodeling, revascularization, and CM dedifferentiation. Overall, these data highlight the regenerative role of *gpnmb* after cardiac cryoinjury in zebrafish.

## 2. Results

### 2.1. *gpnmb* is upregulated in regenerating coronary endothelial cells and in macrophages after cardiac cryoinjury

After cardiac cryoinjury, regenerating coronaries form a scaffold to guide tissue replenishment^12^. We hypothesized that regenerative factors secreted by coronary endothelial cells (cECs) have the potential to direct cardiac repair. To test this hypothesis, we utilized a coronary-specific *Tg(-0*.*8flt1:RFP)* line in combination with a *TgBAC(cxcr4a:YFP)* line to label regenerating cECs after injury. We performed transcriptomic analysis on sorted *-0*.*8flt1:*RFP^+^*cxcr4a:*YFP^+^ cells (regenerating cECs) and *-0*.*8flt1*:RFP^+^*cxcr4a:*YFP^-^ cells (non-regenerating cECs) at 7 dpci (Fig. 1A), a time-point when coronaries extensively cover the injured area ^9,12^. Among the most upregulated genes, we found *gpnmb* strongly induced in regenerating cECs (Fig. 1B). In addition, *cxcr4a* as well as *esm1* and *apln*, which encode endothelial secreted factors known to be involved in angiogenesis, were also upregulated^12,25-28^ (Fig. 1B). Gene Ontology and KEGG analyses of differentially expressed genes (DEGs) (*padj <* 0.05) in non-regenerating vs. regenerating cECs revealed the enrichment of processes related to cardiovascular development and morphogenesis (Fig. S1A). Upregulation of *gpnmb* was further validated by real-time quantitative polymerase chain reaction (RT-qPCR) analyses on zebrafish cryoinjured and sham operated ventricles at 7 dpci (Fig. 1C).

**Figure 1:**
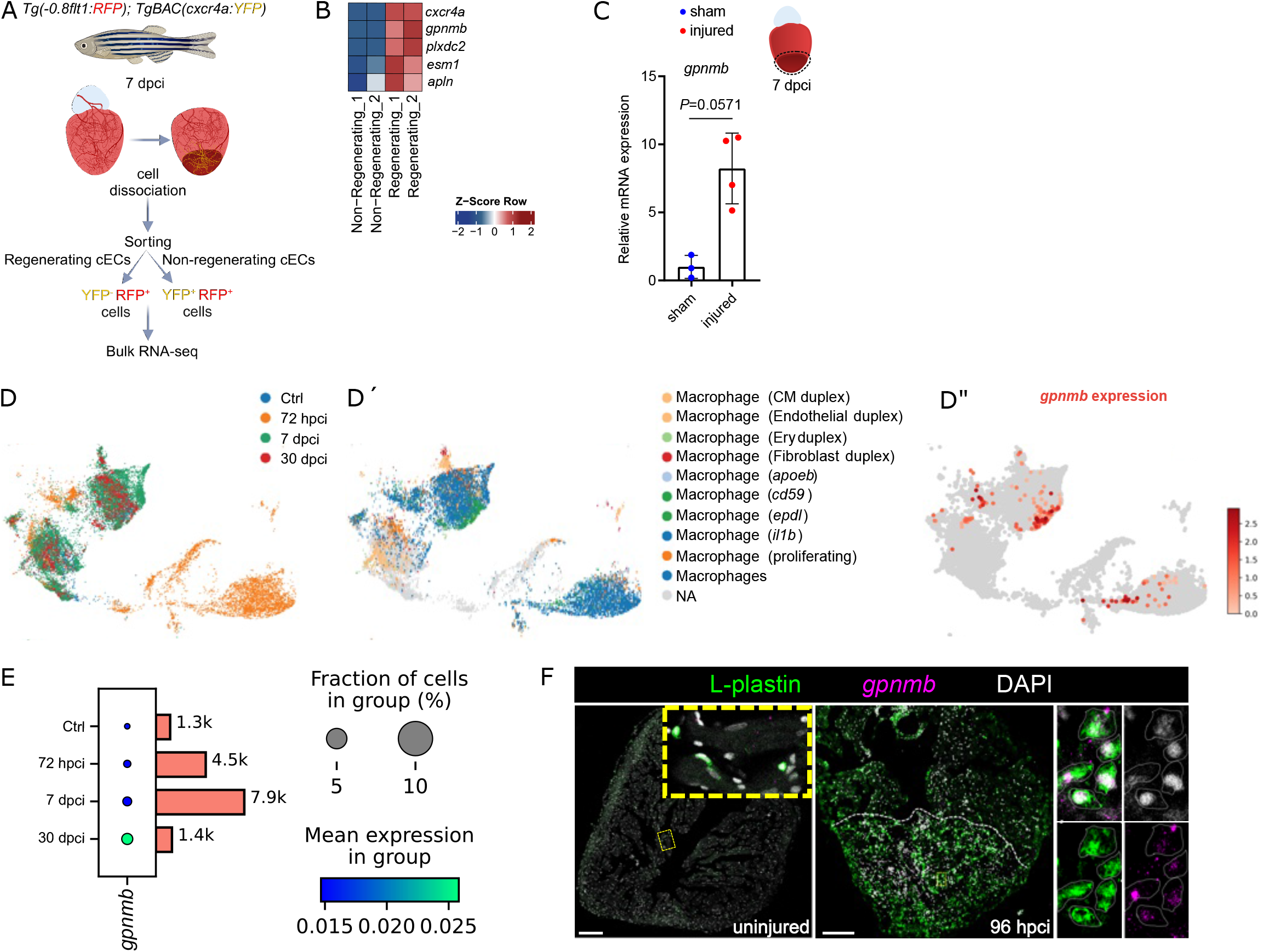
*gpnmb* is upregulated in regenerating coronary endothelial cells and macrophages after cardiac cryoinjury in zebrafish. **A**. Experimental plan for the bulk RNA-seq analysis on sorted regenerating (*-0*.*8 flt1*:RFP^+^/*cxcr4a*:YFP^+^) and non-regenerating (*-0*.*8 flt1*:RFP^+^/*cxcr4a*:YFP^-^) coronary endothelial cells from *Tg(-0*.*8flt1:RFP); TgBAC(cxcr4a:YFP)* whole ventricles at 7 dpci. **B**. Heatmap showing endothelial marker genes (*crcr4a, esm1*, and *apln*) and candidate genes (*gpnmb* and *plxdc2*) in the regenerating and non-regenerating coronary endothelial clusters. **C**. RT-qPCR analysis of *gpnmb* mRNA levels at 7 dpci in the injured tissue relative to sham-operated hearts; Ct values listed in Supplementary Table S1. **D-E**. Reanalysis of the published single-cell RNA sequencing (scRNA-seq) dataset (GSE159032) (Hu et al., *Nature Genetics*, 2022)^29^; UMAP representation of the different macrophage clusters highlighting 1) their distribution at several time-points (D), 2) their identity (D’), and 3) *gpnmb* expression (D’’ ); dot plots of *gpnmb* expression in macrophages (E). **F**. *in situ* hybridization chain reaction (HCR) for *gpnmb* expression and immunostaining for L-plastin on sections of uninjured and 96 hpci ventricles.

In addition, we reanalyzed a published scRNA-seq dataset ^29^ that revealed all the major cardiac cell types (Fig. S1B). We found increased *gpnmb* expression in cryoinjured zebrafish hearts when compared with uninjured control hearts, mainly enriched in certain clusters of macrophages (Fig. 1D-D’’). Subclustering of the macrophages further showed *gpnmb* expression at different stages of regeneration, starting at 72 hpci and with a clear increase at 7 dpci (Fig. 1E) (Fig. S1C). These data are supported by previous reports showing that following cardiac injury in rodents, macrophages are the main cell type expressing *Gpnmb*^24^. To further validate the pattern of expression of *gpnmb* at early time-points after cardiac injury, we performed *in situ* hybridization chain reaction (HCR)^30^ and found *gpnmb* expressed by L-plastin^+^ cells at 96 hpci (Fig. 1F). *gpnmb* expression in sham-operated ventricles was hardly detectable (Fig. 1F). Altogether, these results indicate that macrophages are the main cell type expressing *gpnmb* after cardiac cryoinjury.

### 2.2. *gpnmb* mutants exhibit altered pro- and anti-inflammatory responses

To test the role of *gpnmb* during cardiac regeneration in zebrafish, we generated a *gpnmb* full locus deletion (FLD) allele using CRISPR/Cas9 technology (Fig. 2A). An FLD allele was generated in order to avoid transcriptional adaptation events^31,32^. To assess *gpnmb* expression levels in *gpnmb* FLD mutants, we performed RT-qPCR on 5 days post fertilization (dpf) larvae, adult caudal fins and adult ventricles (injured area) and did not detect *gpnmb* expression in *gpnmb* mutants (Fig. 2A).

**Figure 2:**
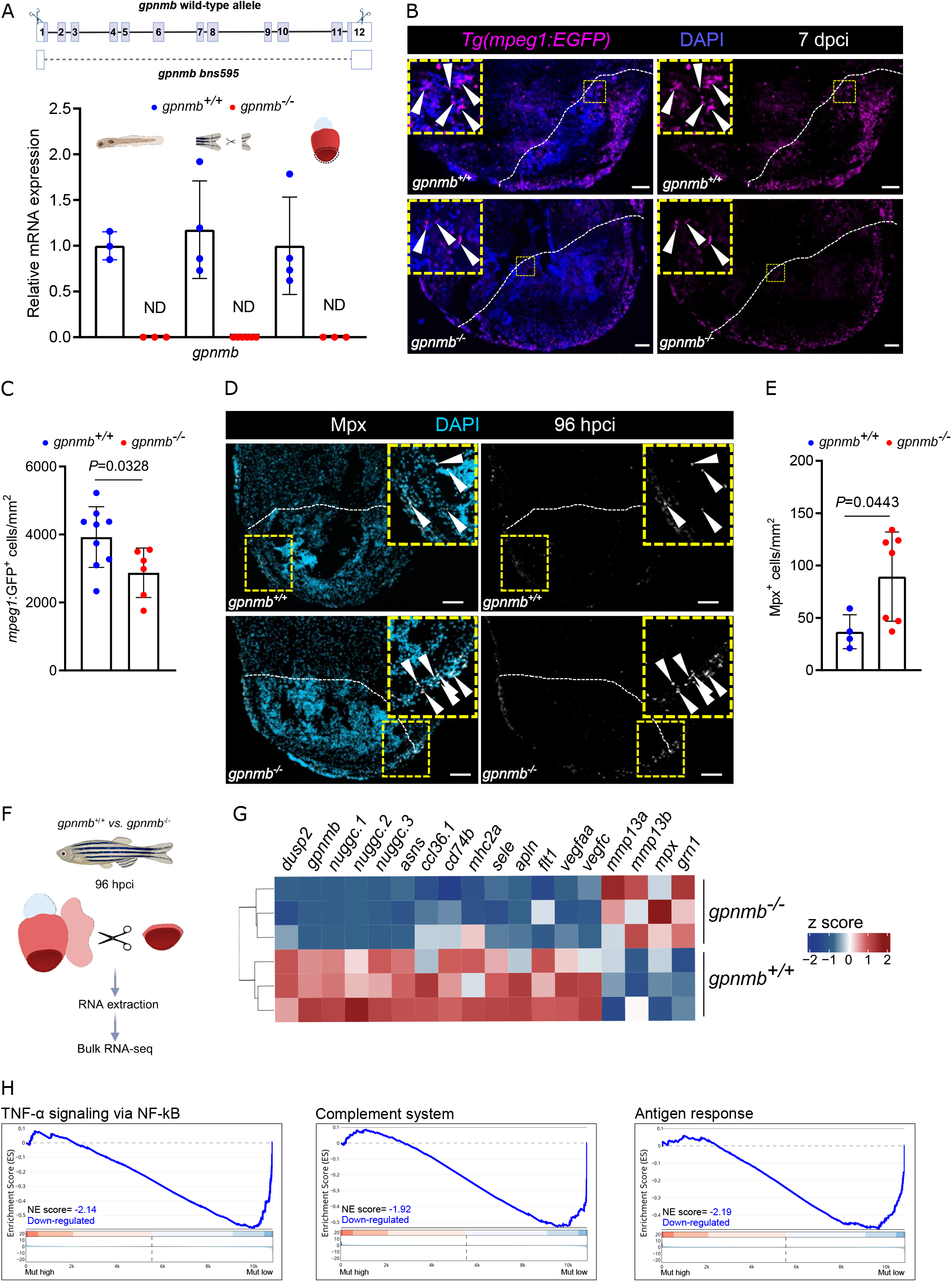
*gpnmb* mutants exhibit altered pro- and anti-inflammatory responses after cardiac cryoinjury in zebrafish. **A**. Illustration of *gpnmb bns595* full locus deletion allele generated using the CRISPR/Cas9 system. RT-qPCR analysis of *gpnmb* mRNA levels in 5 dpf larvae, dissected caudal fins from adult zebrafish, and 7 dpci injured tissue from adult ventricles, comparing *gpnmb*^*+/+*^ and *gpnmb*^*-/-*^ siblings; Ct values listed in Supplementary Table S1. **B**. Immunostaining for EGFP (macrophages, magenta) with DAPI (DNA marker, blue) counterstaining on sections of *Tg(mpeg1:EGFP); gpnmb*^*+/+*^ and *Tg(mpeg1:EGFP); gpnmb*^*-/-*^ cryoinjured ventricles at 7 dpci. **C**. *mpeg1*:EGFP^*+*^ cell number in *gpnmb*^+/+^ and *gpnmb*^-/-^ injured tissues and border zone areas (100 μm) at 7 dpci. **D**. Immunostaining for Mpx (neutrophils, white) with DAPI (DNA marker, blue) counterstaining on sections of *gpnmb*^*+/+*^ and *gpnmb*^*-/-*^ cryoinjured ventricles at 96 hpci. **E**. Mpx^*+*^ cell number in *gpnmb*^+/+^ and *gpnmb*^-/-^ injured tissues at 96 hpci. **F**. Experimental plan for the bulk RNA-seq analysis performed on the border zone and injured (BZI) region of *gpnmb*^*-/-*^ and *gpnmb*^*+/+*^ cryoinjured ventricles at 96 hpci. **G**. Heatmap showing differentially expressed genes of interest in the BZI region of *gpnmb*^*-/-*^ and *gpnmb*^*+/+*^ cryoinjured ventricles at 96 hpci. **H**. GSE analysis plot of TNF-α signaling via NF-kB, complement system and antigen response from *gpnmb*^*-/-*^ vs *gpnmb*^*+/+*^ transcriptomic analysis, 96 hpci. White dotted lines delineate the injured area; white arrowheads point to *mpeg1:*EGFP^+^ (B) and Mpx^+^ (D) cells. Statistical test: Student’s t-test (C, E). Scale bars: 100 μm.

Previous studies indicate that *GPNMB* is highly expressed in macrophages and that it is involved in inflammation resolution and tissue regeneration^17,33^. Other studies have also shown that GPNMB expressing macrophages infiltrate the injured tissue and contribute to injury repair in mouse models^17,21^. Hence, we used the *Tg(mpeg1:EGFP)* macrophage reporter line to assess macrophage presence in cryoinjured *gpnmb*^*-/-*^ ventricles at 96 hpci and 7 dpci, when macrophage numbers gradually increase and peak, respectively^10^. While the number of *mpeg1:*EGFP^+^ cells remained unchanged at 96 hpci (Fig. S2A, B), we found a significant reduction in *mpeg1:*EGFP^+^ cells in *gpnmb*^*-/-*^ ventricles at 7 dpci when compared with wild-type siblings (Fig. 2B, C). Next, to test the dynamics of other leukocyte infiltration^34^, we set out to check whether neutrophil numbers were also affected in *gpnmb* mutants after cardiac injury. Thus, we immunostained cryoinjured *gpnmb*^*-/-*^ ventricles for Mpx, a neutrophil marker, and observed a significant increase in Mpx^+^ cells when compared with *gpnmb*^*+/+*^ ventricles at 96 hpci (Fig. 2D, E). However, no significant difference in neutrophil count was found at 48 hpci (Supplementary Fig. S2C, D).

To gain further mechanistic insight into how *gpnmb* regulates cardiac regeneration, we conducted bulk RNA sequencing on the border zone and injured tissue from cryoinjured *gpnmb*^*+/+*^ and *gpnmb*^*-/-*^ ventricles at 96 hpci (Fig. 2F). We identified 486 differentially expressed genes (DEGs) (*p* < 0.05), with *gpnmb* being one of the most downregulated DEG in *gpnmb*^*-/-*^ vs. *gpnmb*^*+/+*^ (Supplementary Fig. S2E). In line with the phenotype observed, DEG and gene set enrichment (GSE) analyses revealed a decrease in the regulation of immune-related pathways such as TNF-α signaling, the complement system, and the antigen response along with the dysregulation of immune genes including *ccl36*.*1, cd74b, mhc2a, mmp13a, mmp13b, and grn1* (Fig. 2G, H) in cryoinjured *gpnmb*^*-/-*^ ventricles^35,36^. Futhermore, cryoinjured *gpnmb*^*-/-*^ ventricles displayed high levels of genes related to neutrophils (*mpx, csf3r*) coinciding with our results^37^ (Fig. 2D, E, G). Together, these data indicate that *gpnmb* regulates the immune response during heart regeneration.

### 2.3. *gpnmb* mutants exhibit reduced fibroblast activation after cardiac cryoinjury

Immune cells, and specifically macrophages, stimulate fibroblast proliferation and regulate ECM composition by depositing a transient scar^10,38^. The cardiac ECM consists of structural proteins (collagen), adhesive proteins (fibronectins), and de-adhesive tissue-remodeling proteins (tenascins) that play critical roles in the post-infarct zone^39^. Previous studies have suggested the importance of a balanced ratio of collagen, fibronectins, and periostin to facilitate heart regeneration^40^.

GSE (Fig. 3A) and KEGG pathway (Fig. S3A) analyses revealed the enrichment of pathways associated with collagen biosynthesis and assembly, ECM organization, and focal adhesions in cryoinjured *gpnmb*^*-/-*^ ventricles. Our transcriptomic analyses also revealed high expression levels of genes associated with collagens and tenascin (*col10a1a, col11a1a, col12a1b, col11a1b, col17a1b, tnc*) and low expression levels of fibroblast-related genes (*thbs1b, dpt, acta2, postna*) in cryoinjured *gpnmb*^*-/-*^ ventricles^13,35,40^ (Fig. 3B). Previous studies of a mouse liver injury model that lacks Gpnmb-positive macrophages revealed a reduced number of αSMA^+^ cells^21^. αSMA^+^ myofibroblasts are responsible for extracellular matrix remodeling and scar formation after cardiac injury^39^. We analyzed *acta2* expression levels and observed lower levels in injured areas of *gpnmb*^*-/-*^ ventricles at 7 dpci when compared with *gpnmb*^*+/+*^ ventricles (Fig. 3C). To investigate the number of αSMA expressing cells during regeneration in zebrafish, we immunostained for αSMA and observed reduced expression on sections from cryoinjured *gpnmb*^*-/-*^ ventricles at 7 dpci when compared with *gpnmb*^*+/+*^ ventricles (Fig. 3D, E).

**Figure 3:**
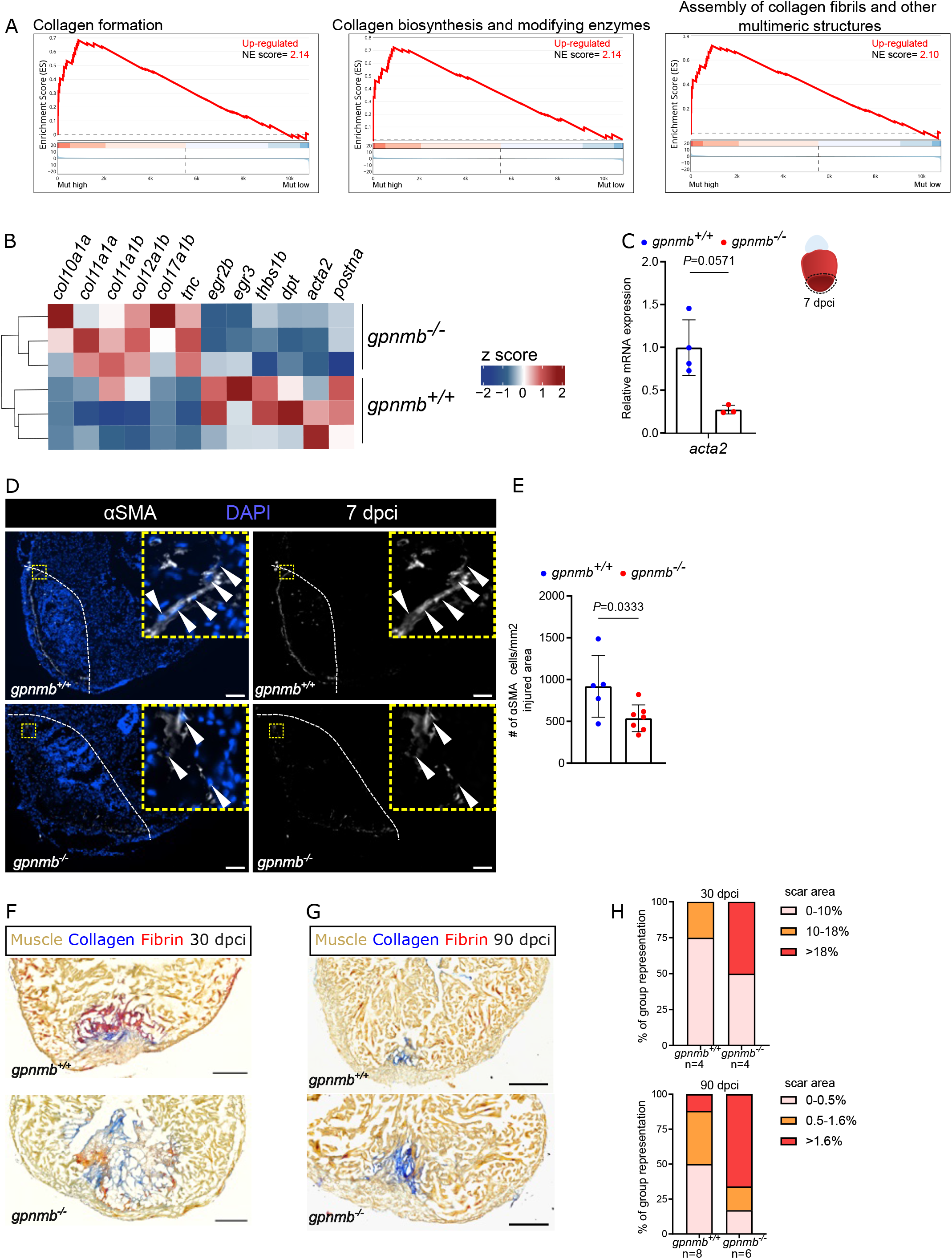
*gpnmb* mutants exhibit reduced fibroblast activation after cardiac cryoinjury in zebrafish. **A**. GSE analysis plot of collagen formation, biosynthesis, and assembly from *gpnmb*^*-/-*^ vs *gpnmb*^*+/+*^ transcriptomic analysis, 96 hpci. **B**. Heatmap showing differentially expressed genes of interest in the BZI region of *gpnmb*^*-/-*^ and *gpnmb*^*+/+*^ cryoinjured ventricles at 96 hpci. **C**. RT-qPCR analysis of *acta2* mRNA levels in the *gpnmb*^*-/-*^ injured tissue normalized to *gpnmb*^*+/+*^ injured tissue at 7 dpci; Ct values listed in Supplementary Table S1. **D**. Immunostaining for αSMA (myofibroblasts, white) with DAPI (DNA marker, blue) counterstaining on sections of *gpnmb*^*+/+*^ and *gpnmb*^*-/-*^ cryoinjured ventricles at 7 dpci. **E**. αSMA^+^ cell numbers in *gpnmb*^+/+^ and *gpnmb*^-/-^ injured tissues at 7 dpci. **F**. AFOG staining on sections of *gpnmb*^*+/+*^ and *gpnmb*^*-/-*^ cryoinjured ventricles at 30 dpci. **G**. AFOG staining on sections of *gpnmb*^*+/+*^ and *gpnmb*^*-/-*^ cryoinjured ventricles at 90 dpci. **H**. Graph showing the representation of groups (y-axis) of different scar area sizes (different colors) at 30 and 90 dpci for *gpnmb*^*+/+*^ and *gpnmb*^*-/-*^ cryoinjured ventricles. Mean of scar area sizes for *gpnmb*^*+/+*^ and *gpnmb*^*-/-*^ at 30 dpci are 9.4 and 11.6 % respectively. Mean of scar area sizes for *gpnmb*^*+/+*^ and *gpnmb*^*-/-*^ at 90 dpci are 0.9 and 1.6 % respectively. White dotted lines delineate the injured area; white arrowheads point to αSMA^+^ cells (E). Statistical tests: Non-parametric Mann-Whitney test (C), Student’s t-test (E). Scale bars: 100 μm (D) and 200 μm (F and H).

As these data indicate that scarring might be affected in *gpnmb*^*-/-*^ mutants, we performed Acid Fuchsin Orange-G (AFOG) staining at 30 dpci when a scar is always present and at 90 dpci when a scar is mostly absent^6^. We found that more *gpnmb*^*-/-*^ ventricles presented larger collagenous scars at 30 and 90 dpci when compared with *gpnmb*^*+/+*^ ventricles (Fig. 3F-H). Altogether, these data suggest that cryoinjured *gpnmb* mutant ventricles exhibit an altered ECM composition and myofibroblast number which might contribute to increased scar size.

### 2.4. Cardiomyocyte and coronary endothelial cell responses are affected in cryoinjured *gpnmb*^***-/-***^ **ventricles**

Timely macrophage recruitment at the injury site is important for neovascularization and cardiomyocyte proliferation, leading to successful heart regeneration in zebrafish^13,34,41,42^. During cardiac regeneration, CMs repopulate the injured region via dedifferentiation and proliferation^11,43,44^. Given the altered inflammatory and fibrotic responses in *gpnmb*^*-/-*^ regenerating hearts along with the impaired function and number of cell types that might impact CM regeneration, we further examined CM dedifferentiation and proliferation. To examine CM dedifferentiation, we immunostained cryoinjured *gpnmb*^*-/-*^ ventricles with the embryonic myosin heavy chain antibody N2.261 which marks dedifferentiating CMs^43,45^. After cryoinjury, *gpnmb*^*-/-*^ ventricles displayed a significant reduction in CM dedifferentiation (Fig. 4A, B) when compared with wild-type siblings. No obvious defects in CM proliferation were observed at 96 hpci, or at 7 or 14 dpci (Fig. 4C, D) (Fig. S3B-E). Next, we set out to test whether *gpnmb* regulates coronary regeneration. We used the *Tg(-0*.*8flt1:RFP)* reporter line, which labels cECs, to assess cEC proliferation in cryoinjured *gpnmb*^*-/-*^ ventricles at 96 hpci, a stage of vigorous cEC proliferation^9,12^ and found a significant increase in cEC proliferation but a decrease in total cEC numbers (Fig. 4E, F) in the injury and border zone areas at 96 hpci. Our transcriptomic analyses also revealed a decrease in the expression of pro-angiogenic genes (*vegfc, vegfaa, flt1*) that are important for the revascularization of the injured tissue post cardiac injury^12,46^. Altogether, these results indicate that the reduced immune response in *gpnmb*^*-/-*^ cryoinjured ventricles affects CM dedifferentiation and cEC number during cardiac regeneration in zebrafish.

**Figure 4:**
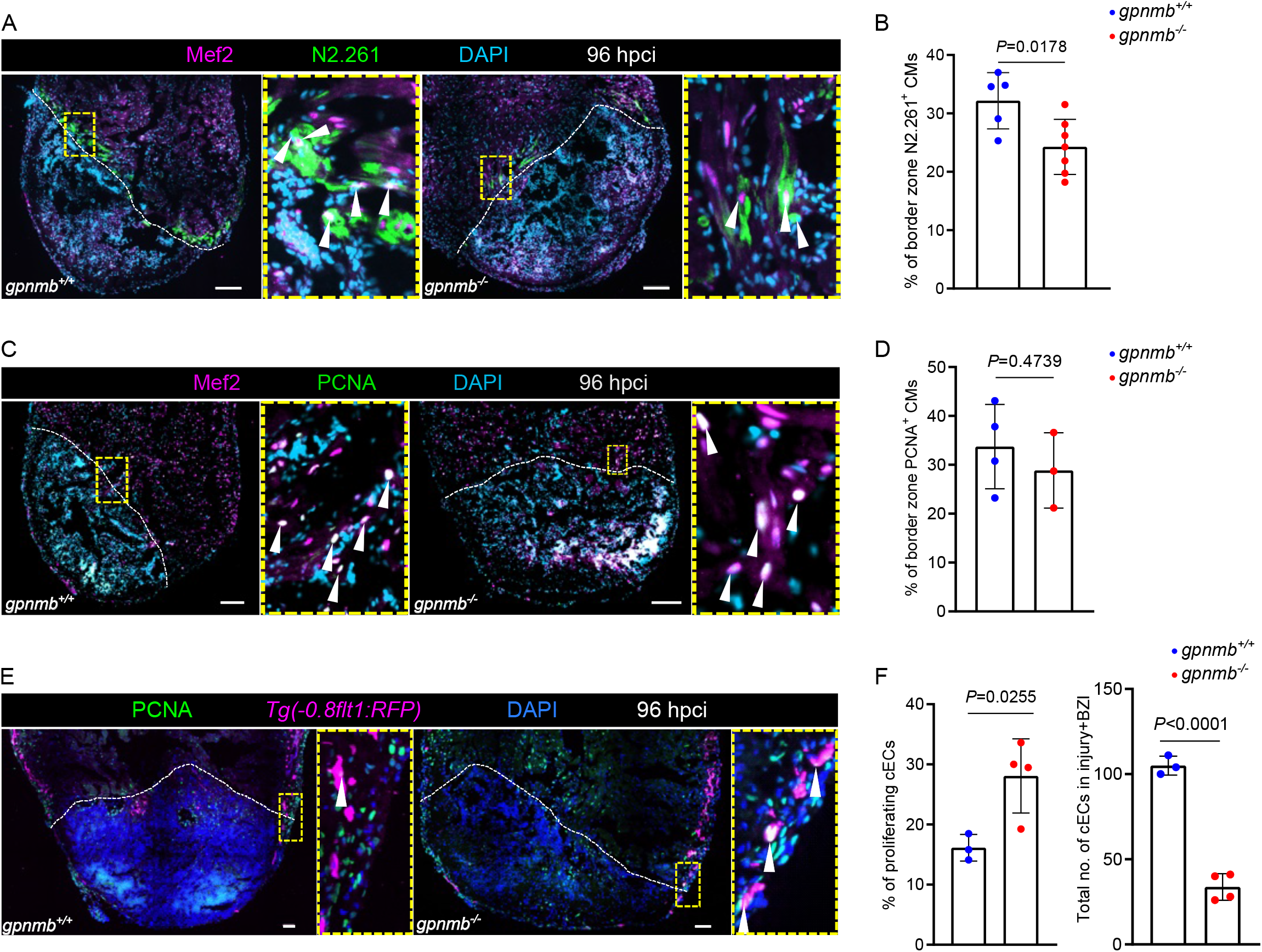
*gpnmb* mutants exhibit increased coronary endothelial cell proliferation and reduced cardiomyocyte dedifferentiation after cardiac cryoinjury in zebrafish. **A**. Immunostaining for MEF2 (cardiomyocyte nuclei, green), N2.261 (embryonic myosin heavy chain, magenta) with DAPI (DNA marker, cyan) counterstaining on sections of *gpnmb*^*+/+*^ and *gpnmb*^*-/-*^ cryoinjured ventricles at 96 hpci. **B**. Quantification of dedifferentiating cardiomyocytes in border zone areas (100 μm) at 96 hpci. **C**. Immunostaining for MEF2 (cardiomyocyte nuclei, green), PCNA (proliferation marker, magenta) with DAPI (DNA marker, cyan) counterstaining on sections of *gpnmb*^*+/+*^ and *gpnmb*^*-/-*^ cryoinjured ventricles at 96 hpci. **D**. Quantification of proliferating cardiomyocytes in border zone areas (100 μm) at 96 hpci. **E**. Immunostaining for PCNA (proliferation marker, green), RFP (coronary endothelial cells, magenta) with DAPI (DNA marker, blue) counterstaining on sections of *Tg(-0*.*8flt1:RFP); gpnmb*^*+/+*^ and *Tg(-0*.*8flt1:RFP); gpnmb*^*-/-*^ cryoinjured ventricles at 96 hpci. **F**. Quantification of proliferating and total number of coronary endothelial cells in injured tissues and border zone areas (200 μm) at 96 hpci. White dotted lines delineate the injured area; white arrowheads point to dedifferentiating cardiomyocytes (A), proliferating cardiomyocytes (C), and proliferating coronary endothelial cells (E). Statistical test: Student’s t-test (B, D, F). Scale bars: 100 μm.

## Discussion

After cardiac injury, temporally controlled tissue inflammation and immune cell recruitment are key to successful regeneration^47,48^. Macrophages, a multifaceted cellular component, are involved in controlling inflammation and are responsible for timely neutrophil clearance, revascularization, CM regeneration, and scar removal after injury^10,13,34,38,41^. GPNMB, a transmembrane glycoprotein, is specifically upregulated in macrophages and other cell types following pro-inflammatory stimuli. While some studies in non-regenerative models suggest a protective role for GPNMB after injury, others have proposed an opposite effect^17,20,23,24,36,49^. Previous studies have showed an increase in GPNMB expression in non-regenerative murine models post myocardial infarction with adverse left ventricular remodeling^24^. In our study, we found that *gpnmb* is required for zebrafish cardiac regeneration as a complete loss of *gpnmb* alters the immune response, reduces myofibroblast numbers, and increases scarring after cardiac injury.

Previous studies in mammals have shown that macrophages upregulate *Gpnmb* in response to tissue damage^20,21,23,24^. In our study, we identified an increase in *gpnmb* expression particularly in macrophages after cardiac injury. The function of GPNMB varies depending on the local microenvironment, cell type, and the interacting ligand. GPNMB overexpression in macrophages plays an active role in the pro- and anti-inflammatory balance; it also promotes the secretion of anti-inflammatory factors and inhibits the secretion of pro-inflammatory factors ^33,50^. After cardiac injury, zebrafish deploy a tightly coordinated inflammatory response with an early pro-inflammatory phase followed by a reparative phase. A balanced immune response is critical to the development and resolution of cardiac fibrosis^13,51^. In line with these findings, we identified neutrophil retention and decreased macrophage recruitment in cryoinjured *gpnmb*^*-/-*^ ventricles. These data support the notion that in the absence of *gpnmb*, inflammation persists in injured mutant hearts, being at least in part the underlying cause of the disrupted regenerative response.

Myofibroblasts modulate the ECM as well as scar deposition. ECM-related processes involve the deposition of a transient collagen-rich scar followed by Tenascin C-associated structural remodeling^3,6,29,40^. Previous studies suggest that GPNMB is a fibroblast activator and GPNMB-treated fibroblasts increase collagen production; however, *Gpnmb* overexpression had no effect on collagen deposition in a denervated muscle model^52^. In addition, reduction in collagen deposition was observed after liver injury in mice lacking *Gpnmb*^21^. However, no quantitative differences were found in collagen deposition in *Gpnmb* WT and mutant mouse hearts after myocardial infarction^24^. Our transcriptomic data show increased collagen expression in cryoinjured *gpnmb*^*-/-*^ ventricles suggesting a role in fibrosis.

Once the soluble form of GPNMB, (*i*.*e*., the extracellular domain) is cleaved in a MMP or ADAM10 protease-dependent manner, it increases vascular density and has a functional role in promoting endothelial cell migration^15^. Revascularization promotes CM replenishment and both processes are essential for cardiac regeneration^9,12,45^. Given the altered inflammatory and fibrotic responses observed in cryoinjured *gpnmb*^*-/-*^ hearts, our findings also suggest that absence of *gpnmb* impairs CM dedifferentiation and cEC numbers.

Overall, our data indicate that *gpnmb* is required for zebrafish cardiac regeneration; it influences inflammation and ECM remodeling, thereby providing important cues for successful regeneration.

## Acknowledgements

We thank R. Ramadass and K. Mattonet for help with microscopy, and P. Goumenaki, T.-L. Tseng and T. Molina-Villa for input during the development of the study and comments on the manuscript. We thank our animal house staff for excellent support. We also thank Philipp Junker (Berlin Institute of Medical Systems Biology) for sharing the pre-analyzed data of GSE159032 for reanalysis. The Marín-Juez lab was supported by the Canadian Institutes of Health Research (PJT-178037). R. Marín-Juez is currently supported by an FRQS Junior-1 award. Research in the Stainier Lab is supported by the Max Planck Society and Leducq Foundation.

## Author Contributions

Conceptualization: S.G., D.Y.R.S and R.M.-J. Methodology: S.G., D.Y.R.S and R.M.-J. Investigation: S.G., G.K., H.E.-S., K.M., S.G., and M.L. Formal analysis: all authors. Writing - Original Draft: S.G., D.Y.R.S and R.M.-J. Writing - Review and Editing: all authors. Supervision: R.M.-J and D.Y.R.S. Project administration: D.Y.R.S and R.M.-J. Funding acquisition: D.Y.R.S and R.M.-J

